# Population genomics of the Anthropocene: urbanization is negatively associated with genome-wide variation in white-footed mouse populations

**DOI:** 10.1101/025007

**Authors:** Jason Munshi-South, Christine P. Zolnik, Stephen E. Harris

## Abstract

Urbanization results in pervasive habitat fragmentation and reduces standing genetic variation through bottlenecks and drift. Loss of genome-wide variation may ultimately reduce the evolutionary potential of animal populations experiencing rapidly changing conditions. In this study, we examined genome-wide variation among 23 white-footed mouse (*Peromyscus leucopus*) populations sampled along an urbanization gradient in the New York City metropolitan area. Genome-wide variation was estimated as a proxy for evolutionary potential using more than 10,000 SNP markers generated by ddRAD-Seq. We found that genome-wide variation is inversely related to urbanization as measured by percent impervious surface cover, and to a lesser extent, human population density. We also report that urbanization results in enhanced genome-wide differentiation between populations in cities. There was no pattern of isolation by distance among these populations, but an isolation by resistance model based on impervious surface significantly explained patterns of genetic differentiation. Isolation by environment modeling also indicated that urban populations deviate much more strongly from global allele frequencies than suburban or rural populations. This study is the first to examine loss of genome-wide SNP variation along an urban-to-rural gradient and quantify urbanization as a driver of population genomic patterns.

## Introduction

Humans exert an outsized influence on ecosystems (Vitousek et al. 1997) in the Anthropocene. This era began sometime between the late Pleistocene (Ellis et al. 2013; Ruddiman et al. 2015) and industrialization in the last few centuries (Steffen et al. 2007), but is always characterized by a global increase in human influence on biological and geochemical processes. Rapid urbanization is a key characteristic of the contemporary Anthropocene. The proportion of humans in cities increased from 16% to 50% in the last century, and is projected to reach 70% by 2050 (Heilig 2011). Urban land conversion may occur at an even faster rate than population growth, thus resulting in accelerating encroachment of cities on reservoirs of biodiversity (Seto et al. 2012). Urban areas become ecologically homogeneous (Groffman et al. 2014) due to loss of vulnerable species that in turn enhances the probability of abrupt state shifts (Barnosky et al. 2012).

Most species are ‘urban avoiders’ that do not persist after urbanization, but ‘urban adapters’ and ‘exploiters’ are facultative or obligate users, respectively, of human-dominated habitats (Blair 2001; McKinney 2002). Although Blair’s (2001) conception of urban adapters did not explicitly include evolution after urbanization, local adaptation may have enhanced the ability of urban adapters to exploit human subsidies. Urban habitats are ‘novel ecosystems’ composed of unique ecological communities and processes (Hobbs et al. 2006; Hobbs et al. 2009), and thus many species likely face new, strong selection pressures from urbanization. Humans have undoubtedly altered the evolutionary trajectory of crop, pest, and disease species (Palumbi 2001), but conclusive cases of human-driven evolution in organisms that are not human commensals or pathogens have been difficult to identify (Merilä and Hendry 2014) outside of a few recent examples (Donihue and Lambert 2014).

Urbanization results in severe habitat fragmentation (Zipperer et al. 2012), a process that reduces genetic variation among animal populations (Keyghobadi 2007; Rivera-Ortíz et al. 2014). Loss of variation may in turn reduce the evolutionary potential of populations experiencing rapidly changing ecological conditions (Etterson and Shaw 2001; Hoffmann and Sgrò 2011; Oakley 2013). Definitive evidence of adaptive evolution from standing variation in wild populations is still relatively scarce (Colosimo et al. 2005; Renaut et al. 2011; Domingues et al. 2012), but new user-friendly approaches to generating large population genomic datasets have improved prospects for quantifying and analyzing genetic variation in the wild (Narum et al. 2013). Evolution can proceed at the nucleotide level through new mutations or changes in frequency of standing genetic variants (Orr 2005), but it is likely that standing variation is more important in cases of rapid evolution (Barrett and Schluter 2008). Results from laboratory experiments (Teotónio et al. 2009; Burke et al. 2010), ancestral variation in model organisms (Rockman 2008; Scarcelli and Kover 2009), humans (Pritchard et al. 2010), artificial selection in crops (Gibson and Dworkin 2004), viral pathogens (Pennings 2012), and invasive species (Prentis et al. 2008) all support this association.

Predicting the loci, genetic architecture, and additive effects involved in adaptive responses to urbanization will be difficult in most cases, but estimating genome-wide variation as a general proxy for evolutionary potential is a useful alternative. Measures of allelic diversity and heterozygosity from genome-wide markers have the advantages of accounting for traits influenced by many loci of small effect, and for predicting how much variation will be available for future adaptive responses to unknown selection pressures (Harrisson et al. 2014). One criticism of using genome-wide SNP datasets to estimate evolutionary potential is that many loci are not associated with functional genomic regions. However, average allele frequency divergence predicts the most extreme *F_ST_* outliers, and the geographic structure of neutral and selected alleles is nearly identical, in humans (Coop et al. 2009). Genome-wide SNPs have also been successfully used to predict phenotypic improvement through “genomic selection” methods in artificially selected species such as livestock (Meuwissen et al. 2013). The increased accessiblity of genome-wide markers for non-model organisms thus provides many opportunities for measuring and predicting evolutionary potential in an urbanizing world (Harrisson et al. 2014). Here, we examine genome-wide variation in white-footed mouse populations sampleed along an urban-to-rural gradient and robustly quantify urbanization as a driver of population genomic patterns.

Previous microsatellite-based analyses on this system showed substantial genetic structure between white-footed mouse populations in NYC’s forest fragments (Munshi-South and Kharchenko 2010). Park area, age, or the extent of habitat within parks did not explain levels of genetic variation among these populations (Munshi-South and Nagy 2014), but these studies focused exclusively on forest fragments within NYC that were highly isolated by surrounding urbanization. In a separate analysis we identified SNPs from transcriptomes sequenced from urban and rural populations, and found that population structure was greater among the urban than the rural populations (Harris et al. 2015). This analysis was limited to only six sampling sites, however, and SNPs from coding regions may often deviate from neutral expectations. In this study we expand these investigations using large, genome-wide SNP datasets, and include populations sampled along an urban-to-rural gradient spanning 142 km from the urban core to extensive, rural protected areas. We specifically used a double-digest RADseq protocol (Peterson et al. 2012) to generate over 10,000 SNPs for analyzing the population genomics of white-footed mice.

Land use transformation and anthropogenic barriers may ultimately reduce gene flow between populations (Epps et al. 2007; Balkenhol and Waits 2009; Jha 2015), leading to greater genetic structuring and loss of genome-wide variation in urbanized areas. Several statistical approaches have recently been developed to investigate the influence of landscapes on genetic structure, although they vary in ability to distinguish between isolation-by-distance (IBD) and ecological factors. Isolation-by-resistance (IBR) models examine the statistical association between genetic and ‘resistance’ distances, where the latter represent probabilities that individuals disperse between populations given the landscape ‘friction’ to dispersal (McRae 2006). The friction values are inferred from empirical movement data or optimized using model selection approaches. One drawback of IBR is that resistance distances are calculated across all paths that individuals may take between populations, and thus are not truly independent from IBD. Isolation-by-environment (IBE) processes, in contrast, are characterized by positive correlations between environmental differences and genetic distances that are independent of the effects of geographic distance (Wang and Bradburd 2014). Both IBR and IBE patterns are influenced by the same biological processes that limit migration across landscapes (such as costs of movement or selection against dispersing genotypes), but IBE patterns are defined by their independence from IBD.

We previously reported that gene flow between populations of white-footed mice could be explained by IBR models based on patterns of urban vegetation cover in NYC (Munshi□South 2012). Here we investigate IBR in a much broader range of landscape conditions along an urban-to-rural gradient. In our earlier work we used partial Mantel tests and a causal modeling approach to identify ecological distances between populations (based on vegetation cover) that best explained gene flow after factoring out IBD (Cushman and Landguth 2010). While this approach can be successful for landscapes composed of high-contrast cover types (such as NYC), several authors have argued that partial Mantel tests have low statistical power and are prone to false positives (Legendre and Fortin 2010; Graves et al. 2013). Here we use a partial Mantel test for our IBR model, as well as a new statistical IBE approach to quantify the relative contributions of urbanization and IBD to genetic differentiation between white-footed mouse populations (G. S. Bradburd et al. 2013). Specifically, we model the strength of covariance in allele frequencies between populations as a function of IBD and IBE due to urbanization of the landscape.

In this study, we test the following interrelated predictions about the population genomics of white-footed mice (*Peromyscus leucopus* Rafinesque) in the New York City (NYC) metropolitan area:

1. Evolutionary potential as measured by genome-wide variation within populations is inversely related to urbanization of the surrounding landscape.
2. Urbanization of the landscape results in greater genomic structure and differentiation between populations.
3. “Resistance distances” and “ecological distances” resulting from urbanization are better predictors of genetic differentiation than geographic distances between populations.

## Methods

### Study species and sampling sites

White-footed mice are one of the most widespread and abundant small mammals in eastern North America, and occupy a broad range of forest, meadow, and secondary growth habitats. They have served as model systems for population ecology for decades (Vessey and Vessey 2007; Brunner et al. 2013) because of their ubiquity and easy trappability. *Peromyscus* spp. more broadly have emerged as model systems for the genomics of adaptation (Bedford and Hoekstra 2015), and the first reference genomes are currently being assembled (Kenney-Hunt et al. 2014).

For this study, we sampled white-footed mice from 23 sites in the NYC metropolitan area (Table 1). The 12 sites within NYC limits were the same as in previous studies, although here we combined samples from Pelham Bay and Hunters Island in the Bronx, and Alley Pond and Cunningham Parks in Queens, because evolutionary clustering results showed that these pairs of sites were not strongly differentiated from each other (Munshi-South and Kharchenko 2010). These urban sites contained secondary forest typically dominated by oaks, hickories, maples, and/or tulip trees, with very thick understories composed primarily of invasive plants. Suburban and rural sites contained similar canopies but the understories were largely cleared of thick vegetation by rampant deer herbivory.

**Table 1.**
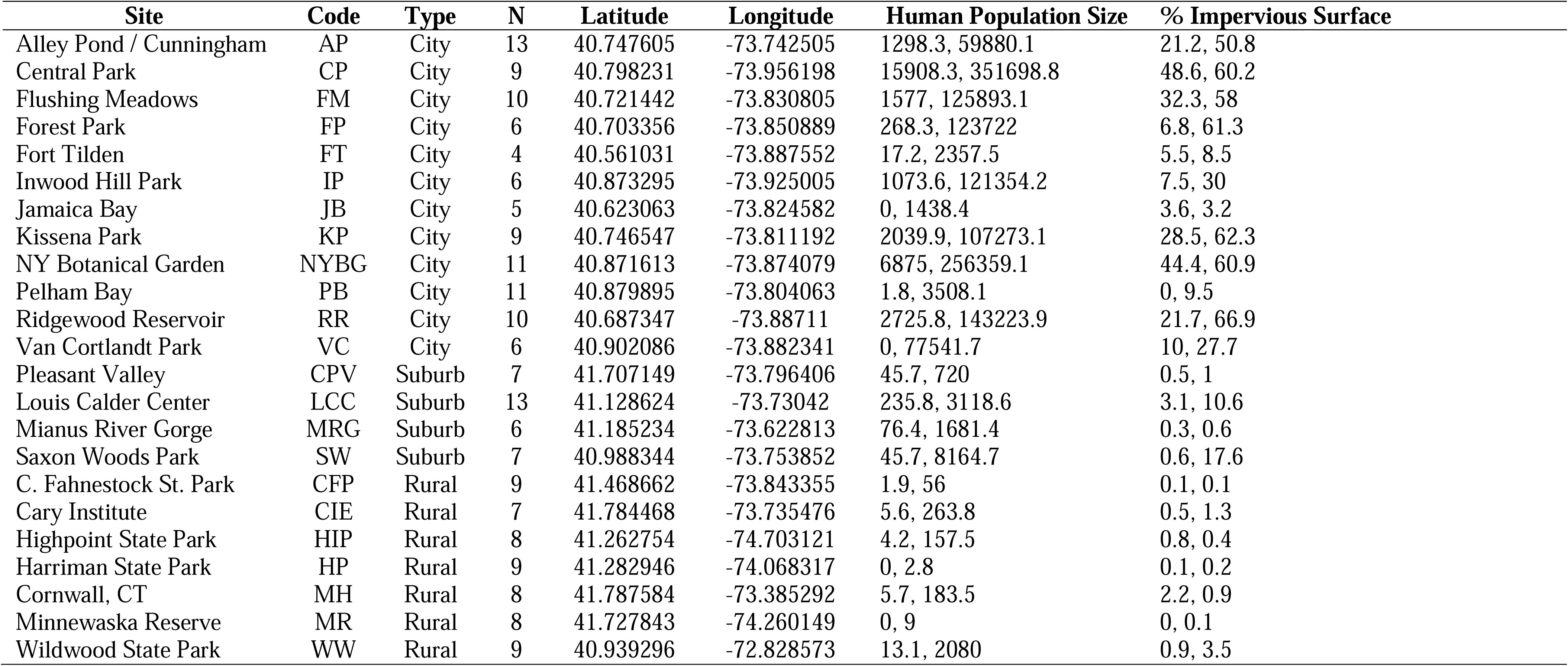
Characteristics of numbers of white-footed mice sampled at 23 locations along an urban-to-rural gradient. Human population size and percent impervious surface were measured at two buffer sizes around these sites: 500 m and 2.0km.

| Site | Code | Type | N | Latitude | Longitude | Human Population Size | % Impervious Surface |
| --- | --- | --- | --- | --- | --- | --- | --- |
| Alley Pond / Cunningham | AP | City | 13 | 40.747605 | -73.742505 | 1298.3, 59880.1 | 21.2, 50.8 |
| Central Park | CP | City | 9 | 40.798231 | -73.956198 | 15908.3, 351698.8 | 48.6, 60.2 |
| Flushing Meadows | FM | City | 10 | 40.721442 | -73.830805 | 1577, 125893.1 | 32.3, 58 |
| Forest Park | FP | City | 6 | 40.703356 | -73.850889 | 268.3, 123722 | 6.8, 61.3 |
| Fort Tilden | FT | City | 4 | 40.561031 | -73.887552 | 17.2, 2357.5 | 5.5, 8.5 |
| Inwood Hill Park | IP | City | 6 | 40.873295 | -73.925005 | 1073.6, 121354.2 | 7.5, 30 |
| Jamaica Bay | JB | City | 5 | 40.623063 | -73.824582 | 0, 1438.4 | 3.6, 3.2 |
| Kissena Park | KP | City | 9 | 40.746547 | -73.811192 | 2039.9, 107273.1 | 28.5, 62.3 |
| NY Botanical Garden | NYBG | City | 11 | 40.871613 | -73.874079 | 6875, 256359.1 | 44.4, 60.9 |
| Pelham Bay | PB | City | 11 | 40.879895 | -73.804063 | 1.8, 3508.1 | 0, 9.5 |
| Ridgewood Reservoir | RR | City | 10 | 40.687347 | -73.88711 | 2725.8, 143223.9 | 21.7, 66.9 |
| Van Cortlandt Park | VC | City | 6 | 40.902086 | -73.882341 | 0, 77541.7 | 10, 27.7 |
| Pleasant Valley | CPV | Suburb | 7 | 41.707149 | -73.796406 | 45.7, 720 | 0.5, 1 |
| Louis Calder Center | LCC | Suburb | 13 | 41.128624 | -73.73042 | 235.8, 3118.6 | 3.1, 10.6 |
| Mianus River Gorge | MRG | Suburb | 6 | 41.185234 | -73.622813 | 76.4, 1681.4 | 0.3, 0.6 |
| Saxon Woods Park | SW | Suburb | 7 | 40.988344 | -73.753852 | 45.7, 8164.7 | 0.6, 17.6 |
| C. Fahnestock St. Park | CFP | Rural | 9 | 41.468662 | -73.843355 | 1.9, 56 | 0.1, 0.1 |
| Cary Institute | CIE | Rural | 7 | 41.784468 | -73.735476 | 5.6, 263.8 | 0.5, 1.3 |
| Highpoint State Park | HIP | Rural | 8 | 41.262754 | -74.703121 | 4.2, 157.5 | 0.8, 0.4 |
| Harriman State Park | HP | Rural | 9 | 41.282946 | -74.068317 | 0, 2.8 | 0.1, 0.2 |
| Cornwall, CT | MH | Rural | 8 | 41.787584 | -73.385292 | 5.7, 183.5 | 2.2, 0.9 |
| Minnewaska Reserve | MR | Rural | 8 | 41.727843 | -74.260149 | 0, 9 | 0, 0.1 |
| Wildwood State Park | WW | Rural | 9 | 40.939296 | -72.828573 | 13.1, 2080 | 0.9, 3.5 |

At each site, we trapped white-footed mice over a period of one to three nights using two or more 7x7 grids of Sherman live traps (9”x3”x3”). Traps within grids were placed 15 m apart, and grids were located several hundred meters apart to avoid trapping close relatives. We collected ear punches or tail clippings from up to 25 mice at each site, and stored the tissue in 80% EtOH before transfer to a -20 C freezer in the laboratory. Animal handling procedures were approved by Fordham University’s Institutional Animal Care and Use Committee (Protocol No. JMS-13-03).

We chose sites that qualitatively represented typical urban (*N* = 12 sites), suburban (*N* = 4), and rural (*N* = 7) sites in the NYC metropolitan area based on levels of development (Figure 1). However, suburban counties adjacent to NYC have higher human population densities than major cities in other parts of North America. To facilitate comparisons with other urban areas, we calculated the size of the human population and percent impervious surface in geographic buffers around each site as proxies for relative urbanization (Figure 1b).

**Figure 1.**
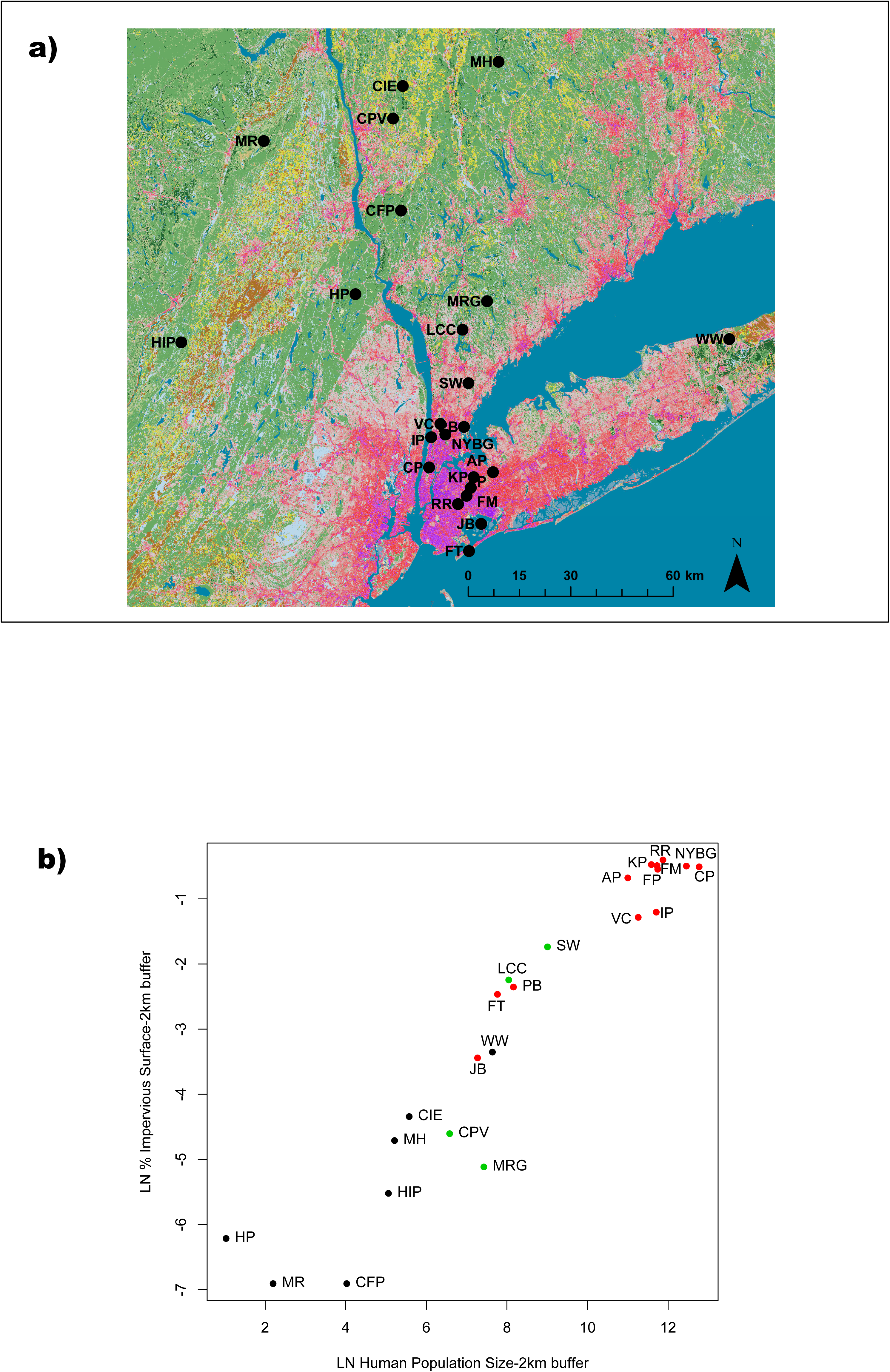
**a.** Geographic locations of the 23 sites at which we sampled white-footed mice for this study. The map contains land cover categories and impervious surface levels from the 2011 National Land Cover Database at 30 m resolution. Red to purple colors represent increasing percentages of impervious surface cover. Site abbreviations correspond to Table 1. **b.** Scatterplot of percent impervious surface cover vs. human population size for the 23 sampling sites. Both variables were log-transformed to improve linearity, and were measured using 2 km buffers around study sites as described in the text. Red dots represent urban sites, green dots represent suburban sites, and black dots represent rural sites.

We first incorporated GPS coordinates for our study sites into base layers (2010 TIGER/Line^®^ Shapefiles—County or States and equivalent) available from the US Census Bureau. Buffers were created in ArcGIS v10.1 (ESRI, Redlands, CA) around each site’s GPS coordinates with radii of 500 m, 1,000 m, 1,500 m and 2,000 m. These buffers were chosen because they are relevant to the typical lifetime dispersal distance of many white-footed mice; for example, no mice over a 40 year study of one woodlot dispersed to the nearest neighboring woodlot less than 1.5 km away (Vessey and Vessey 2007). To determine human population size inside each buffer, we used U.S. Census Blocks as these provide the smallest geographic unit with 100% census data. We first calculated the area of each census block and then intersected the census block and buffer layers. We interpolated human population size within each buffer based on the percentage of area for each census block that intercepted each buffer ([area of census block within buffer / area of census block] x population of census block). We then summed all of the interpolated population sizes that fell within each buffer.

To determine percent impervious surface within each study site buffer, we used the 2011 Percent Developed Imperviousness layer from USGS National Land Cover Data (Xian et al. 2011). We then calculated zonal statistics to measure the geometry of the raster file and summarize the cell values of the raster that fell within each buffer. From these data we calculated the average percent impervious surface within each landscape buffer for each site.

These estimates of impervious surface and human population size could be highly correlated at the different spatial buffers, so we calculated Pearson correlation coefficients between all pairs of values (Table S1). To avoid including redundant information in downstream analyses, we removed any variables that exhibited *r* > 0.90 in pairwise comparisons with the other variables. The 1 km and 1.5 km buffers were removed because they were highly correlated with other estimates, and percent impervious surface and human population size estimated for the 500 m and 2 km buffers were retained.

### ddRADSeq, SNP genotyping, and population genomic statistics

We generated SNP genotypes for 233 individuals using a double-digest RADseq protocol (Peterson et al. 2012). We aimed to obtain genotypes from 10 individuals from each site, but sample dropout due to poor DNA yield during library preparation resulted in variability in the sample size for each site (Table 1). In brief, we extracted DNA from tail snips or ear punches using Qiagen DNEasy tissue kits (Qiagen, Valencia, CA) with an RNAse treatment, and then digested 500-1,000 ng of DNA using the enzyme combination of SphI-HF and MluCI for one hour following the manufacturer’s instructions (New England Biolabs, Ipswich, MA). A Qubit 2.0 fluorometer (Life Technologies, Norwalk, CT) was used to quantify DNA concentration at several steps in the library preparation. Next, digested DNA was cleaned with 1.5X AMPure XP magnetic beads (Beckman-Coulter, Brea, CA) and custom in-line ‘flex’ barcodes and P1/P2 adapters were ligated to 200-400 ng of digested DNA for each sample (Peterson et al. 2012). Up to 48 individual libraries with unique barcodes were then pooled in equimolar amounts and cleaned with 1.5X AMPure XP beads. Next we selected DNA fragments of known sizes (376-412 bp) from the pooled libraries using a Pippin Prep (Sage Science, Beverly, MA). We then conducted multiple PCR amplifications using 20 ng of size-selected DNA and Phusion High-fidelity PCR reagents with manufacturers’ PCR conditions (New England Biolabs, Ipswich, MA). This PCR step added a second, unique index sequence and Illumina sequencing primers to the pooled libraries so each individual sample contained a unique combination of the in-line barcode and index. We then pooled the PCRs for each size-selected library, cleaned the pools using 1.5X AMPure XP beads, and checked the libraries using an Agilent BioAnalyzer (Agilent Technologies, Santa Clara, CA) for DNA concentration and the correct distribution of fragment size. The libraries were sequenced using three lanes of Illumina HiSeq 2000 2x100 bp paired-end sequencing at the New York University Center for Genomics and Systems Biology (New York, NY, USA).

As an initial check on the quality of our Illumina sequence data, we analyzed the raw reads in FastQC (Andrews 2010). Subsequent demultiplexing, quality filtering and *de novo* SNP calling was conducted using the Stacks 1.21 pipeline (Catchen et al. 2013). First, we used the *process_radtags* script to filter out low-quality reads and demultiplex the remaining reads according to their unique combination of in-line barcode and index. Based on FastQC results, we trimmed all reads to 96 bp to remove poor-quality base calls at the ends of reads. Next, we concatenated the single-and paired-end reads for each individual into one fastq file because the two paired reads did not overlap and were not aligned to a reference genome. To identify RAD loci and call SNPs, we used the *denovo_map.pl* script in Stacks with default settings except for the minimum number of identical reads required to create a ‘stack’ (m = 7), number of mismatches allowed between RAD loci for a single individual (M = 3), and the number of mismatches allowed when building the catalog (n = 2). After building the initial catalog of loci, we used the *populations* script in Stacks to filter loci for those that occurred in at least 22 / 23 sampling sites (p = 22) and in at least 50% of individuals at each site (r = 0.5). This script also produced genotype output in multiple formats (i.e. Genepop, Structure) with one randomly-selected SNP from each locus (--write_random_SNP), and generated summary statistics such as observed heterozygosity, nucleotide diversity, and pairwise FST between all populations (Table 2).

**Table 2.**
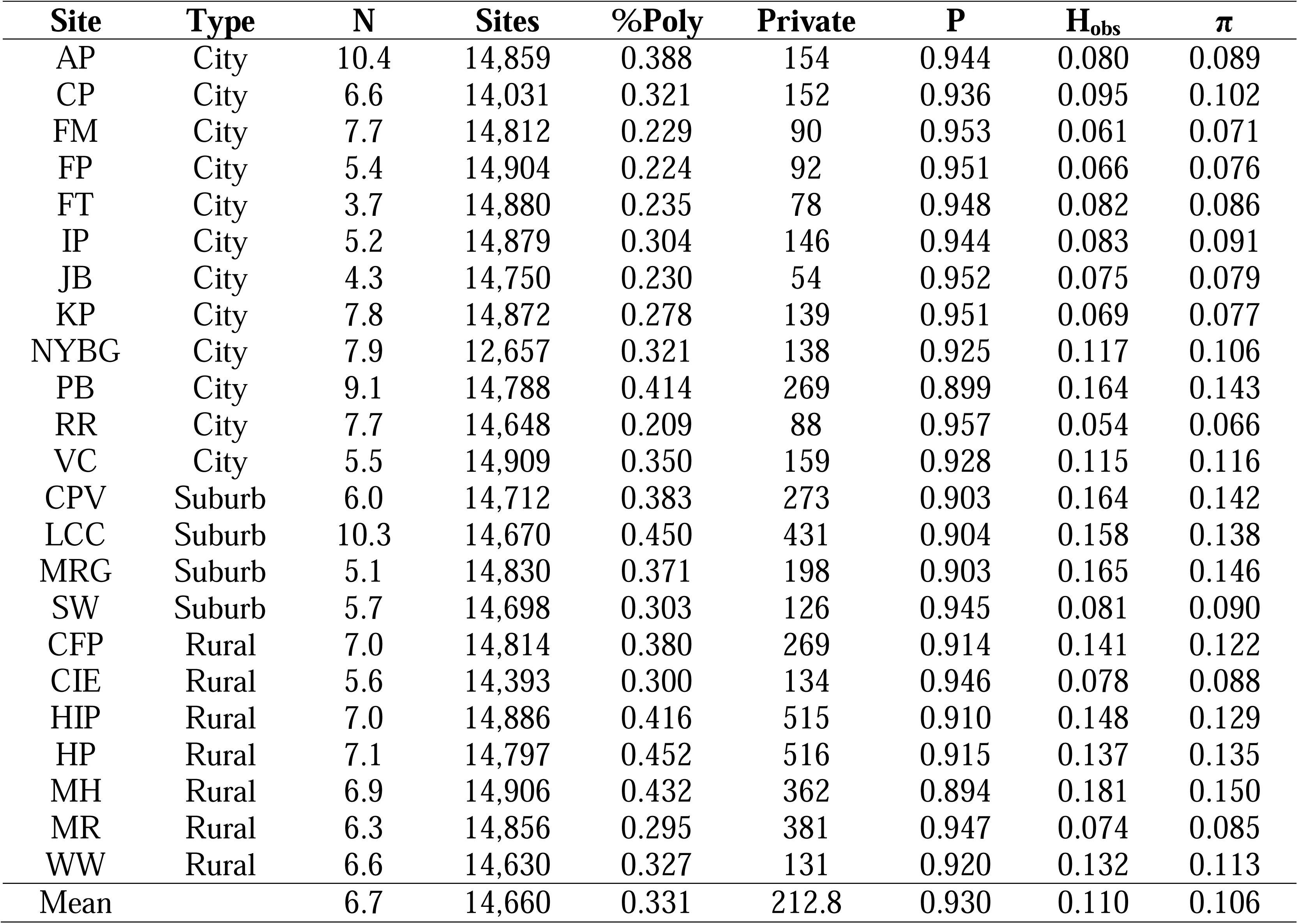
Summary genetic diversity statistics calculated by STACKS for nucleotide positions that were polymorphic in at least one population. *N* = average number of individuals genotyped at each locus; *Sites* = number of polymorphic nucleotide sites across the dataset; *%Poly*= percentage of polymorphic loci; *Private* = number of variable sites unique to each population; *P* = average frequency of the major allele; *H_obs_* = average observed heterozygosity per locus; *π* = average nucleotide diversity.

We loaded the RAD loci and individual data from Stacks into a MySQL database and visualized the output using the Stacks webserver. Based on the results of the first pipeline run, we removed 26 individuals because their small number of reads resulted in very small SNP datasets and excessive missing genotypes compared to other samples. Highly related individuals in our dataset could also bias downstream analyses. We avoided relatives at our urban sites using relatedness values from previous microsatellite studies of the same individuals (Munshi-South and Kharchenko 2010). For the SNP dataset, we identified highly related individuals by calculating kinship coefficients in the software package KING (Manichaikul et al. 2010).

Kinship analysis identified 16 pairs of individuals that were related at the half-sib level or greater. We removed one of the pair members from the dataset, resulting in a final dataset comprised of 191 of the original 233 individuals. We then re-ran the Stacks pipeline on the screened dataset of 191 individuals.

Besides the filters applied by Stacks and the removal of highly-related individuals, we also omitted outlier loci detected using the approach in BayeScan 2.1 (Foll and Gaggiotti 2008). Operating with default parameters and a False Discovery Rate < 0.01, we identified 200 outliers using BayeScan. Five of these outliers showed a signature of positive selection, whereas 195 showed signatures of balancing selection. We omitted these outliers from the dataset because they did not fit assumptions of neutrality inherent to our downstream analyses.

### Modeling genome-wide variation within populations

We modeled four measures of genome-wide variation within populations against our two urbanization proxies to test the hypothesis that urbanization is associated with reduced genetic variation. Summary statistics for each population included observed heterozygosity (*Ho*), nucleotide diversity (π), number of private loci, and the percent of polymorphic loci. We examined genetic (dependent) variables using seven candidate general linear models (GLMs) consisting of combinations of human population size and percent impervious surface estimated using different geographic buffers as described above (500 m and 2,000 m): an intercept-only model, four univariate models with one variable measured at one buffer size, and two bivariate models including both human population and impervious surface estimated for the same buffer size. Human population size and percent impervious surface were ln-transformed prior to analysis to improve normality. If a model performed substantially better than the intercept-only model, then we interpreted that result as evidence of a statistical effect of the urbanization variable(s) on genetic diversity within white-footed mouse populations. We calculated maximum likelihood estimates of model parameters for each model, and then ranked models using values of *AICc,* the corrected Akaike’s information criterion (Burnham and Anderson 2002). The relative quality of models was further assessed based on Δ*AICc*, and the relative weight (*w_i_*) of each model. For the best GLMs, we examined the statistical significance of the urbanization coefficients. We also examined scatterplots (Figure 2) and fitted regression lines to confirm that the model explained variation in the genetic parameter of interest. We conducted all statistical analyses in R 3.2.1 (R Development Core Team 2008), and used GLM regression in the AICcmodavg package for model selection (Mazerolle 2015).

**Figure 2.**
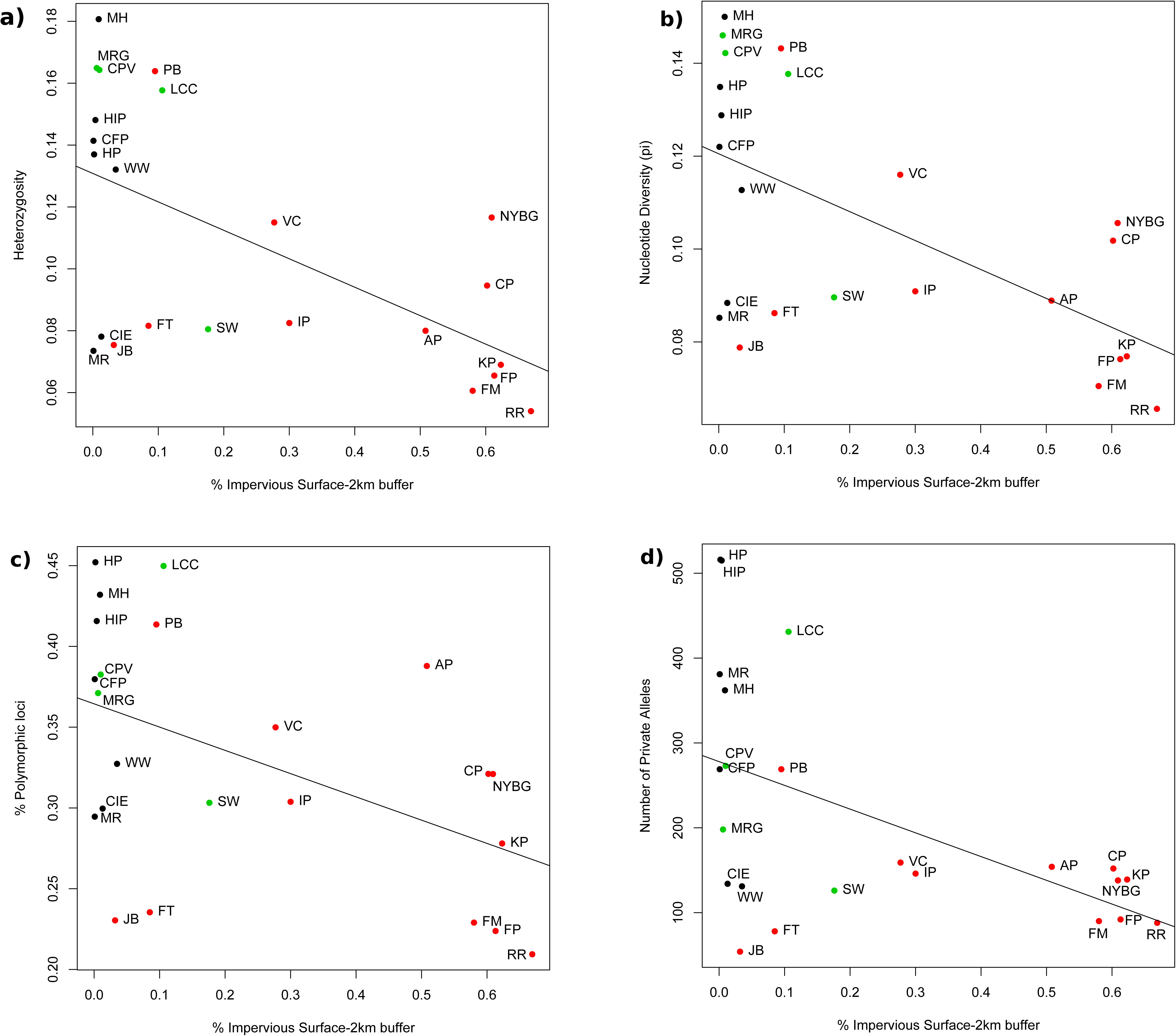
Scatterplots and trend lines for best GLMs identified using AIC_c_ modeling that describe the relationship between urbanization and **a)** heterozygosity, **b)** nucleotide diversity, **c)** percent polymorphic loci, and **d)** number of private alleles for the 23 populations. Red dots represent urban sites, green dots represent suburban sites, and black dots represent rural sites. Site abbreviations correspond to Table 1.

### Genetic structure and population differentiation

To investigate whether urbanization results in greater genetic differentiation between populations, we used Discriminant Analysis of Principal Components (DAPC) in the R package *adegenet* (Jombart and Ahmed 2011) to identify evolutionary clusters among 191 white-footed mice. DAPC first reduces total genetic variation (i.e. variance in allele frequencies) into principal components, and then identifies discriminant functions that maximize differences between clusters while minimizing variation within clusters. We used cross-validation in *adegenet* to identify the optimal number of principal components. This procedure uses a randomly-generated training set and validation set of individuals to identify the optimal number of principal components that accurately predict group membership without overfitting. To visualize clusters, we used DAPC scatterplots and barplots (compoplot command in *adegenet*) to visualize membership of individuals in different clusters. Genome-wide SNP datasets often have power to discriminate between all groups, and results from analyses such as DAPC may reflect hierarchical structure. Thus, we ran DAPC on the full dataset as well as subsets of sampling sites to investigate hierarchical structure.

We also used the model-based evolutionary clustering approaches in fastSTRUCTURE (Raj et al. 2014) and ADMIXTURE (Alexander et al. 2009). fastSTRUCTURE uses approximate inference of the Bayesian model in the original STRUCTURE (J. Pritchard et al. 2000) whereas ADMIXTURE computes maximum likelihood estimates of parameters to estimate the most likely number of evolutionary clusters, K. We ran fastSTRUCTURE on our data for each value of K from 1-23 using the standard model with a simple prior. A fastSTRUCTURE script (*chooseK.py*) also calculate heuristic scores for detecting the range of the most likely values of K. After identifying likely values of K, we re-ran fastSTRUCTURE with the more computationally demanding logistic prior model on a smaller subset of K values. The most likely values of K were determined in ADMIXTURE using its cross validation procedure (Alexander and Lange 2011). To compare clustering results from fastSTRUCTURE and ADMIXTURE, we used the CLUMPAK (Cluster Markov Packager Across K) web server (Kopelman et al. 2015) to align and visualize bar plots for both programs at multiple values of K.

### Landscape genomics

To investigate IBD, IBR and IBE between white-footed mouse populations, we first tested for IBD using a Mantel test in the *ecodist* package in R. The data were bootstrapped 10,000 times to generate 95% confidence intervals for the Mantel *P* value. We then used the BEDASSLE package (Bradburd 2014) in R to estimate the relative contributions of IBD and IBE to genetic differentiation between the sampling sites. We computed allele counts and sample sizes for each population using the --counts function in VCFtools 0.1.12b (Danecek et al. 2011). We examined the following IBE models using BEDASSLE: 1) a simple binary matrix indicating whether the population was located in NYC or outside the city; and 2) a pairwise matrix of “resistance distances” calculated between populations using the IBR approach in Circuitscape 4.0 (McRae and Beier 2007).

Circuitscape calculates a pairwise matrix of resistance distances between populations based on the ability of a simulated electrical current to flow between adjacent landscape cells connected by resistors with user-defined resistance values. We used the 2011 Percent Impervious Surface layer from USGS National Landcover Data (Xian et al. 2011), and set the resistance value for each 30 m cell as equivalent to its percent impervious surface unless the cell exceeded 70% impervious surface. For cells exceeding 70%, we set the resistance level to 100; in other words, any cell with greater than 70% impervious surface was assumed to be 100X more resistant to migration than a cell with 1% impervious surface. The 70% cutoff for relatively high, quasi-barrier resistance was based on results of an earlier IBR analysis we conducted in NYC (Munshi□South 2012). To calculate resistance distances between all populations, we ran Circuitscape in pairwise mode with raster cells connected to all eight neighboring cells. The analysis also produced a cumulative current map to visualize hypothesized migration between all populations. As an initial check on the success of the resistance distances at explaining variation in pairwise F_ST_, we conducted a partial Mantel test with 10,000 bootstraps in *ecodist* that factored out the effects of Euclidean geographic distance. We then used these resistance distances for the IBE model in BEDASSLE.

BEDASSLE uses a Markov Chain Monte Carlo (MCMC) approach, and includes several graphing functions for evaluating success of the MCMC posterior parameter estimation. To confirm adequate mixing and convergence of chains, we examined traces and marginal distributions for all parameters. We also examined acceptance rates for MCMC parameter estimation, and when these rates were too high or low we adjusted the tuning parameters and reran the analysis for 5-10 million steps. For the final runs, we calculated the median and 95% credible intervals for the E: D ratio after discarding the first 20% as burn-in. This ratio represents the effect size of ecological distance relative to geographic distance.

The simple BEDASSLE model assumes identical variance of allele frequencies about the global mean allele frequency. However, populations deviate from the global mean for a number of demographic reasons (i.e. bottlenecks, inbreeding), and outlier populations can have a strong influence on posterior distributions. To account for this variation, BEDASSLE includes a beta-binomial model that estimates an additional parameter, Φ_K_, that measures the strength of drift and lack of fit of each population to the model. To address population history, we also ran the beta-binomial model for the two IBE scenarios above. For these models, we examined the posterior distributions of the Φ_K_ values (recalculated as F_K_ = 1 / 1+ Φ_K_) to identify outlier populations.

To test the relative fit of the models to our data, we generated 1,000 posterior predictive samples for each model and compared the simulated data to our observed data in BEDASSLE. BEDASSLE includes a function that randomly draws parameter values from the posterior distributions and simulates new datasets. These simulated datasets are then used to calculate F_ST_ between all pairs of populations. Simulated F_ST_ values can then be compared to the observed F_ST_ values to examine the models’ ability to describe patterns in the real data.

## Results

### ddRADSeq dataset and summary population genomic statistics

Three lanes of Illumina HiSeq 2x100 PE sequencing produced a total of 1.16 billion reads, of which 936.64 million reads (80.9%) passed initial quality filters. Reads were filtered out due to ambiguous barcode sequences (56.3%), low quality scores (40.9%), and ambiguous RAD tags (2.8%). The catalog generated by STACKS included 880,898 RAD loci with at least 7X coverage, but this number was reduced to 14,930 after requiring loci to be present in 22 / 23 populations and at least 50% of individuals within each population. All populations were well-represented in the final dataset, with averages of 4 to 10 individuals genotyped per locus across the 23 populations (median = 6.6; Table 2).

For all loci that were polymorphic in at least one population, the major allele frequency (P) ranged from 0.894 to 0.957, and the average observed heterozygosity from 0.054 to 0.181 (Table 2). If all invariant positions are included, then the major allele frequency exceeded 0.999 in all populations and the heterozygosity ranged only from 0.001 to 0.002 (Table S2). Nearly all of the populations located within NYC exhibited lower genetic variation than suburban or rural populations as measured by the percentage of polymorphic sites, numbers of private alleles, major allele frequency, observed heterozygosity, and nucleotide diversity (π). The only exceptions to these trends were relatively high diversity values for the Pelham Bay (PB) population in NYC, and low values for suburban Saxon Woods (SW), the rural Cary Institute (CIE) and Minnewaska Reserve (MR; Table 2). Excluding those four outliers, heterozygosity ranged from 0.054 to 0.117 in the urban populations and 0.158 to 0.181 in the suburban and rural populations. Nucleotide diversity similarly ranged from 0.066 to 0.116 in NYC and 0.138 to 0.15 in the suburban and rural populations.

### Modeling urbanization and genome-wide variation within populations

Sampling sites could be distinguished by their combinations of human population size and percent impervious surface (Figure 1b). The rural sites all clustered at very low values for both urbanization variables. Urban and rural sites exhibited little overlap in the scatterplot, with the exception of Fort Tilden (FT), Jamaica Bay (JB), and Pelham Bay in NYC. Figure 1b shows the relationship between the two urbanization variables measured for a 2 km buffer around sampling sites, but results were qualitatively similar for the 500 m buffer.

Model selection based on *AIC_c_* confirmed that nearly all GLMs with one or more urbanization variables described variation in genomic diversity better than intercept-only models, except for human population size estimated at a 500 m buffer (Table 3). Impervious surface cover estimated at the 2 km buffer,was the highest-ranked model for all four genetic variables, with the second best models all including both impervious surface cover and human population size at the 2 km buffer. The Akaike weights for the best models were all equal to or greater than 0.55, and the delta *AICc* for moving from the second to third model were all greater than 3.0. Heterozygosity was negatively associated with percent impervious surface, with the exception of the outlier populations mentioned above (i.e. rural populations with low genetic diversity; Figure 2a). Nucleotide diversity was also negatively associated with percent impervious surface, and exhibited the same outlier populations as the heterozygosity scatterplot (Figure 2b). The number of private alleles and percent polymorphic loci were negatively associated with percent impervious surface (Figure 2c,d). All urbanization coefficients from the top GLMs were also significant at *P* < 0.05, with the coefficients for nucleotide diversity and number of private alleles significant at *P* < 0.001.

**Table 3.**
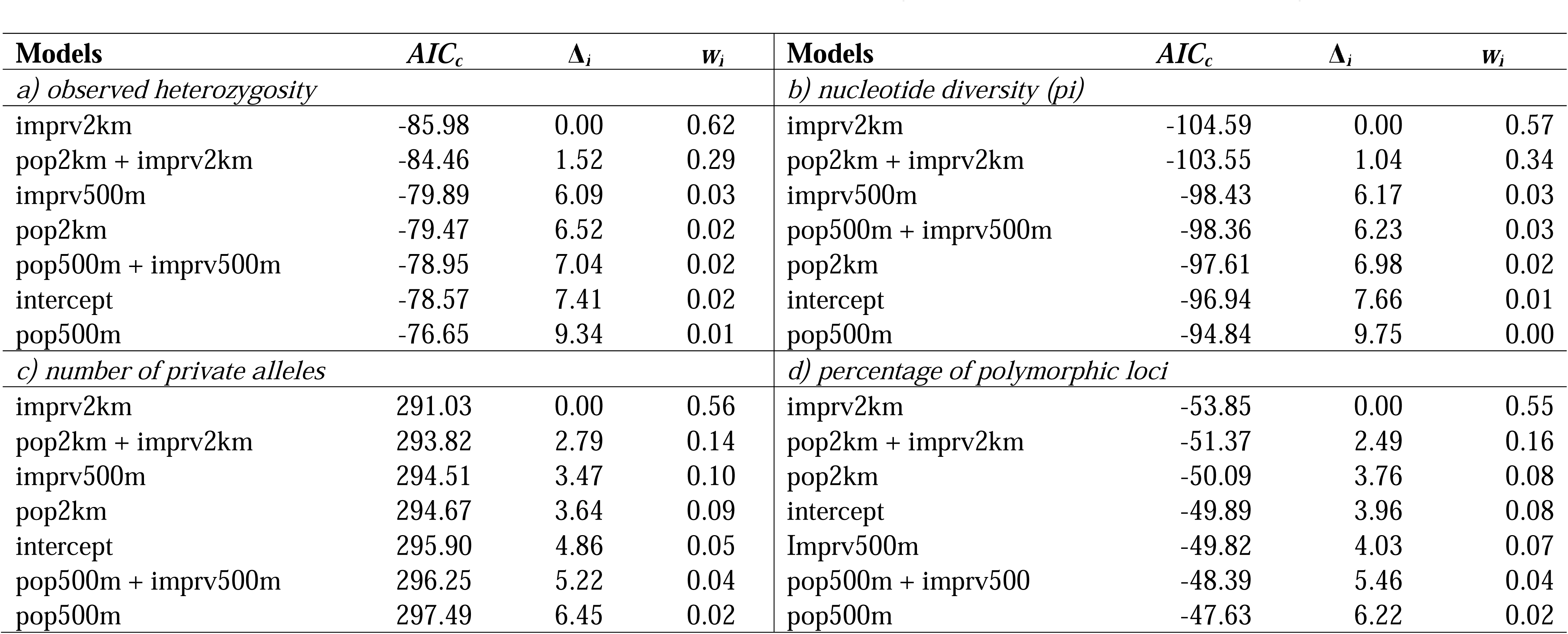
Results of selection among general linear models describing the influence of percent impervious surface cover (imprv) and human population size (pop) at 500 m and 2 km buffer sizes on a) observed heterozygosity, b) nucleotide diversity, c) the number of private alleles within each population, and d) the percentage of polymorphic loci within each population. The best models were chosen based on the second-order Akaike Information Criterion (*AIC_c_*), the rate of change in AIC_c_ (Δ*_i_*) and the Akaike weights (*w_i_*).

### Genetic structure and population differentiation

Pairwise *F_ST_* calculated using STACKS ranged from a low of 0.033 between a rural (MH) and suburban (LCC) population, and a high of 0.145 between a suburban (MRG) and urban (RR) population. Most values were between 0.05 – 0.10. Some NYC populations exhibited many *F_ST_* > 0.10, particularly RR and JB, as did one suburban population (MRG). A simple Mantel test revealed no significant IBD (Mantel *r* = 0.041, 95% CI = -0.046 – 0.124, *P* = 0.32).

Cross-validation identified *N* = 23 as the optimal number of principal components to retain for DAPC analysis (Figure S1). The first two discriminant functions distinguished two isolated NYC sites, Jamaica Bay (JB) and Fort Tilden (FT), from the other populations. Discriminant function three separates out a cluster of the other populations on Long Island (AP, FM, FP, KP, RR, WW), a cluster of NYC populations in the middle (IP, NYBG, PB, VC) with suburban populations just below them (LCC, MRG, SW, CPV), and rural populations (CIE, CFP, HIP, HP, MH, MR) at the bottom (Figure 3a). This clustering largely recapitulates the spatial orientation of these populations along a north-south axis. One exception is the most urban population, Central Park (CP), which does not occur where it would be expected based on geography. This placement indicates that Central Park is one of the most isolated and unique NYC populations, along with Fort Tilden and Jamaica Bay. The DAPC compoplot (i.e. a barplot of membership probability) indicated that nearly all individuals could be assigned to their sampling site with high probability (Figure S2).

**Figure 3.**
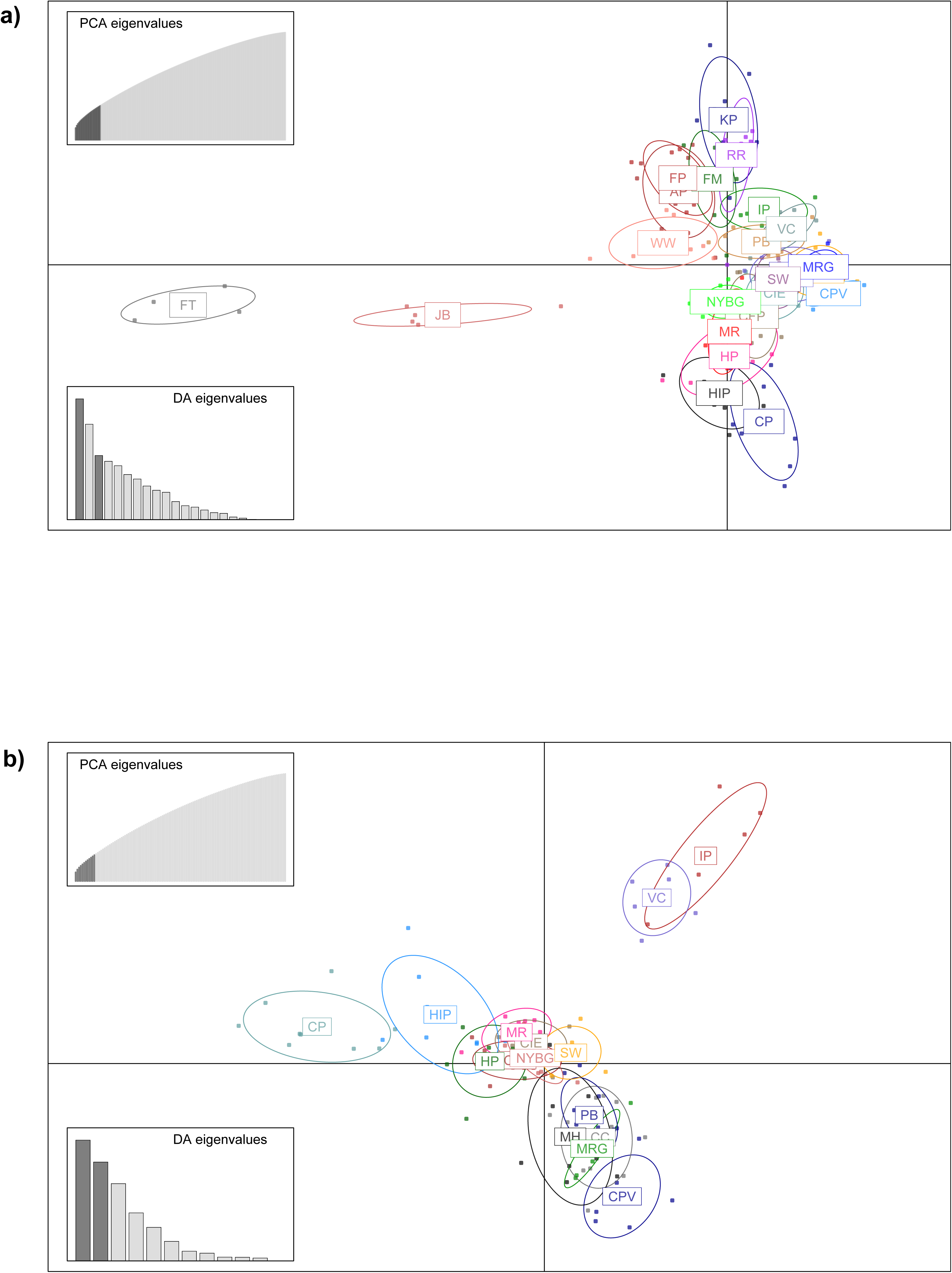
Scatterplots resulting from Discriminant Analysis of Principal Components (DAPC) for **a)** all 23 sampling sites, and **b)** a subset of sites occurring on Manhattan and mainland North America. Insets represent the eigenvalues of retained principal components (top left), and the eigenvalues of discriminant functions portrayed in the scatterplots (bottom left). Site abbreviations correspond to Table 1.

To clarify relationships between major clusters of sampling sites, we reran the DAPC analysis on two subsets: all populations on the mainland and Manhattan, and all populations located on Long Island. Twelve principal components were retained for the mainland-Manhattan analysis, and the scatterplot of the first two discriminant functions revealed a major, central cluster that recapitulated the geography of the populations. However, three isolated, urban populations were distinct from this cluster (Figure 3b): Central Park (CP), Inwood Hill Park (IP), and Van Cortland Park (VC). The Long Island analysis retained seven principal components, and confirmed that JB and FT are highly distinct populations, as well as the Ridgewood Reservoir (RR; Figure S3).

Both the heuristic analysis in fastSTRUCTURE (using the simple prior) and cross-validation in ADMIXTURE identified K = 2 as the most likely number of evolutionary clusters among the 23 white-footed mouse populations we sampled. CLUMPAK confirmed that individual assignment to the two clusters was highly correlated across the two methods (*r* = 0.93; Figure 4a,b). One cluster (blue in Figure 4a,b) contained all the NYC populations on Long Island, the two populations on Manhattan (CP and IP), and three other suburban / rural populations with atypically low genetic variation (CIE, MR, and SW). All other populations were assigned to the other cluster (orange in Figure 4a,b), except for WW located in rural Long Island which was an admixture of the two clusters in almost equal proportions. When we reran the fastSTRUCTURE analysis for K=1 – 4 using the more accurate logistic prior, the heuristic analysis identified K = 3 as the upper bound on the likely number of evolutionary clusters. Cross-validation in ADMIXTURE also identified K = 3 – 5 as only slightly worse than K = 2 (Figure S4), so we created barplots for these numbers of evolutionary clusters. For K = 3, the blue cluster of urban Long Island, Manhattan, and outlier rural populations was maintained, but the other cluster was split into two largely based on location east or west of a north-south axis (Figure 4c). The fourth cluster in the K = 4 ADMIXTURE analysis included a new cluster (Figure 4d) with the two Manhattan populations, a Bronx population (NYBG), and a few distant populations adjacent to or west of the Hudson River (CFP and HIP). The additional cluster for K = 5 included two suburban populations in relative proximity (CPV and LCC; Figure 4e).

**Figure 4.**
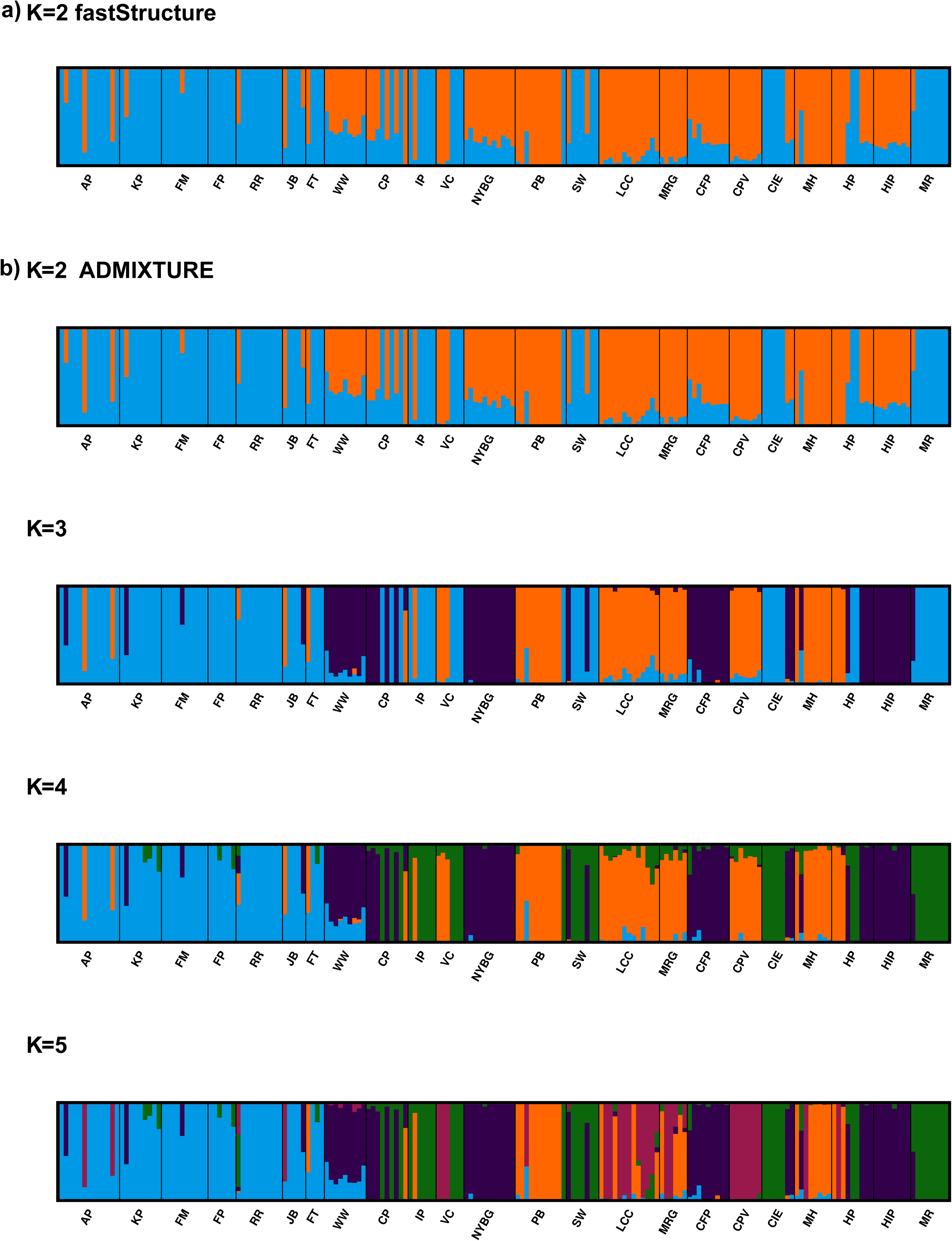
Bar plots resulting from evolutionary clustering analyses using a) fastSTRUCTURE assuming K = 2, and b) using ADMIXTURE assuming K= 2 – 5. Site abbreviations correspond to Table 1, and are ordered geographically in the bar plots. The first five sites are on Long Island (AP-WW), the next two on Manhattan (CP and IP), the next 10 east of the Hudson River (VC-MH, organized from North to South), and the last three west of the Hudson River (HP-MR).

### Landscape genomics

For the simple BEDASSLE model examining presence or absence in the city, the median α_E_: α_D_ ratio was 7.14 and the 95% credible set was 6.90 to 7.36. The interpretation of this result is that being located in NYC has an impact of approximately 7 km of extra pairwise geographic distance on genetic differentiation. Comparison of posterior predictive samples for the simple and beta binomial model confirms that the latter is a much better fit to the data (Figure 5a,b). For the beta binomial model, the median α_E_: α_D_ ratio was 0.006 and the 95% credible set was 0.0002 to 0.038. This model indicates that presence in NYC has virtually no additional impact over geographic distance on genetic differentiation. However, values of *F_k_* were elevated for most of the urban populations (range in medians from 0.04 – 0.25; mean = 0.12) relative to the suburban (range = 0.01 – 0.06; mean = 0.03) and rural populations (range = 0.001 – 0.11; mean = 0.03), with the exception of two rural populations that also had high values (CIE and MR; Figure 5d). These *F_k_* values indicate that the urban populations deviated substantially more from global allele frequency estimates than the suburban and rural populations.

**Figure 5.**
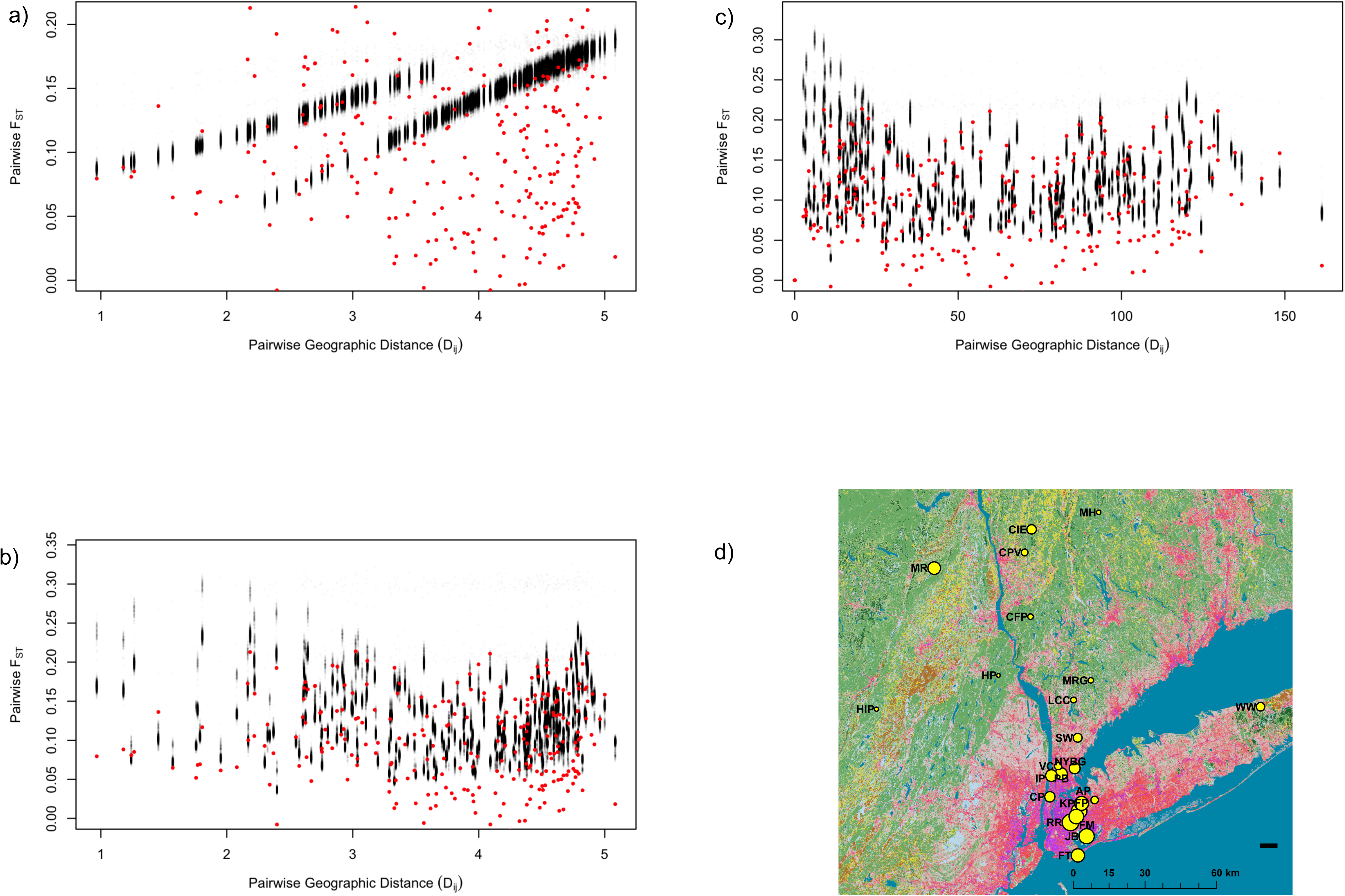
Posterior predictive sampling with 1,000 simulated datasets in BEDASSLE, using pairwise F_ST_ as a summary statistic for a) the simple IBE model with presence in or outside NYC as the environmental variable; b) the beta binomial IBE model for presence in or outside NYC; and c) the beta binomial IBE model with resistance distances based on percent impervious surface as the environmental variable. d) Map of study sites with points scaled using the values of *F_K_* estimated by the beta binomial model in BEDASSLE. *F_K_* estimates the deviation of each population from global mean allele frequencies, and was highest for populations located within NYC.

Pairwise resistance distances from the IBR model (Figure S5) were significantly associated with F_ST_, even after factoring out the effects of geographic distance (partial Mantel *r* = 0.521, 95% confidence interval = 0.438-0.621, *P* < 0.0001; Table S3). The simple IBE model using resistance distances did not converge after trying many different combinations of prior values. The beta binomial model did converge, producing a median α_E_: α_D_ ratio of 0.0001 and 95% negligible impact on genetic differentiation compared to geographic distances, which contradicts the IBR results. Posterior predictive sampling indicated that this beta-binomial model performed moderately well, but systematically underpredicted observed F_ST_ < 0.05 (Fig. 5c).

## Discussion

This study is the first to our knowledge to examine the influence of urbanization on genome-wide SNP variation in a city-dwelling species. We found that populations of white-footed mice along an urban-to-rural gradient exhibit a negative correlation between genomic variation and urbanization in and around their habitat. Populations within NYC also exhibited greater genetic differentiation from one another than pairs of populations in rural areas. IBR and IBE models based on urbanization explained a greater proportion of pairwise population differentiation overall than IBD by some metrics. NYC populations deviated more strongly from global mean allele frequencies than rural populations, indicating that urbanization has substantially altered the evolutionary trajectories of urban wildlife (Donihue and Lambert 2014).

Many recent studies have documented reduced migration and loss of heterozygosity at microsatellite loci among populations in urbanized landscapes (Gortat et al. 2014; Jha 2015; Barr et al. 2015). However, for logistical reasons the most variable microsatellite loci are often chosen for population genetic analysis, and thus may not represent unbiased samples of genome-wide diversity (Väli et al. 2008). Heterozygosity measured using microsatellites and traits related to fitness are also often weakly correlated, even though there is a publication bias towards reporting only high heterozygosity-fitness correlations (Chapman et al. 2009). New approaches such as ddRAD-Seq used here produce genome-wide SNP markers that are more appropriate for assessing genome-wide variation in relation to ecological factors such as urbanization.

Population genomic variation is key to understanding and predicting evolutionary responses to environmental transformation. While urbanization may not directly cause extinctions of most species, it is likely to decrease the evolutionary potential of populations. Results from native species in cities are few. Urban bobcats maintained variation at immune-linked loci due to balancing selection from disease pressures, even after a population bottleneck and population subdivision by freeways (Serieys et al. 2015). Blackbirds colonizing cities also exhibited polymorphisms in a candidate behavioral gene that was strongly associated with urban habitats (Mueller et al. 2013), indicating that functional variation is important for the evolutionary success of urban wildlife. We previously identified candidate genes that may be under selection in urban white-footed mice (Harris et al. 2013), but our results here indicate that these populations have lost as much as half of their genetic diversity compared to nearby rural populations. While candidate genes of large effect may be relevant to understanding some responses to landscape change, screening genome-wide variation offers many advantages for measuring evolutionary potential. Most adaptive traits are polygenic and influenced by many loci of small effect. Information on genome-wide variation will also capture cryptic variation and quantify the amount of standing variation available for responses to future change (Harrisson et al. 2014). Loss of standing variation will thus make it less likely that urban populations will be able to adapt to local conditions, or to global phenomena such as climate change (Franks et al. 2014). Mutation is likely too slow of a process for rapid recovery of evolutionary potential in fragmented urban populations. However, increasing connectivity in cities would immediately boost genetic variation if migration between genetically differentiated populations could be reestablished.

### Modeling urbanization and genome-wide diversity

We found that percent impervious surface cover, and to a lesser extent human population size, estimated at 2 km buffers around study sites were highly correlated with levels of genome-wide diversity. Gradient studies have predominated in urban ecology, but terms such as “urban” and “suburban” have been used in many different contexts without standardization (Magle et al. 2012). Study sites are often defined based on subjective criteria, or chosen simply to represent a linear geographical gradient regardless of the actual pattern of urbanization (Ramalho and Hobbs 2012). We advocate that future landscape genetics studies report percent impervious surface cover in and around study sites to facilitate comparisons between landscapes and species. Impervious surface and human population size are highly correlated, but humans may be present in large numbers outside of cities for recreation in protected areas (Monz et al. 2013). Impervious surface can be readily used to track urbanization over time, and has relevance to both terrestrial and aquatic systems (Walsh et al. 2005). Extensive impervious surface in the form of roads, parking lots, and buildings characterizes all cities (Nowak and Greenfield 2012). Roads in particular have well-characterized negative impacts on the connectivity of wildlife populations (Balkenhol and Waits 2009; Benítez-López et al. 2010). Nearly 70% of global forest cover is now fragmented, and areas subject to urbanization and high-intensity agriculture are most severely affected (Haddad et al. 2015). Reporting measures of impervious surface cover will provide much needed standardization. We found that estimates for 2 km buffers were much better than 500 m buffers, although the spatial effects will likely vary for different species. In this case, 500 m was likely too small of a buffer to capture the extent of urbanization’s influence on study sites, whereas 2 km was near the maximum buffer size we could use around many of our study sites and still retain statistical independence from other study sites.

### Population structure and differentiation

Urban and rural populations were differentiated from each other, with many *F_ST_* > 0.10. Some populations within NYC had pairwise *F_ST_* as high as urban-rural pairs that were much more distant, indicating that isolation within the city is quite high. Higher *F_ST_* values reported here are similar to those reported for endangered beach mice (Austin et al. 2015) and Channel Island deer mice (Ozer et al. 2011) that recently experienced strong genetic drift from extirpations and translocations. The lack of IBD reported here, but moderate to strong genetic population differentiation, suggests that dispersal is limited by barriers and high landscape resistance rather than geographic distance. Thus, evolutionary clustering and IBE (Wang and Bradburd 2014; Sexton et al. 2014) are more appropriate models for understanding patterns of genome-wide diversity among these populations.

Discriminant analysis of principal components, and two model-based clustering analyses, sorted individuals into 23 clusters that largely recapitulated geographic patterns. We identified one major split between Long Island and mainland populations, which historical demographic modeling indicated is likely related to glacial retreat and ecological succession in the region (Harris et al, unpublished manuscript). The other striking pattern was that several urban populations were outliers in their genetic divergence from the major clusters of sampling sites. Previous clustering analyses using microsatellites also identified most NYC sites as distinct populations (Munshi-South and Kharchenko 2010). SNPs evolve more slowly than microsatellites, and there is likely still considerable ancestral variation in isolated urban populations. These NYC populations may also not have reached linkage and Hardy-Weinberg equilibrium at many SNP loci because not enough generations have elapsed since isolation, or the populations have not reached mutation-migration-drift equilibrium.

### Landscape genomics

Our IBR and IBE analyses produced mixed results. Overall, IBR and IBE modeling indicated that urbanization drives genetic differentiation to a greater degree than geographic distance alone. Although the relative influence of IBE to IBD was modest in the BEDASSLE analyses, urban populations deviated to a much greater degree from global allele frequencies than suburban or rural populations. The most likely explanation for this deviation is substantial drift due to inbreeding or bottlenecks in isolated urban habitats. This scenario generally conforms to the strong structure observed among some of the urban populations. However, other factors that caused these populations to deviate from the BEDASSLE model cannot be ruled out, such as unsampled environmental variables (Bradburd et al. 2013).

IBR modeling in Circuitscape confirmed that variation in percent impervious surface is highly associated with variation in pairwise *F_ST_*, even after factoring out IBD. Partial Mantel tests have several known issues with false positives (Legendre and Fortin 2010), but we previously found that vegetation cover (typically the inverse of impervious surface in NYC) successfully described migration between populations using similar approaches (Munshi□South 2012). The IBE models did not identify the same clear association between impervious surface and genetic differentiation. As above, the strong deviation of urban populations from the global mean allele frequencies indicated that the IBE model may have underperformed. Resistance distances from Circuitscape are integrated over all possible paths between points on the landscape, and thus BEDASSLE also counts geographic distance twice with unknown consequences when using resistance distances (G. Bradburd, personal communication).

IBE is an active area of inquiry that holds great promise for understanding the processes that generate genetic variation. However, currently available approaches are relatively new, and several caveats apply to their application (Wang and Bradburd 2014). This study is the first use of BEDASSLE to model the relationship between genetic differentiation and human modification of the environment. In addition to the model adequacy issues raised above, some of the populations analyzed here may not have reached migration-drift equilibrium, or were dominated by idiosyncratic environmental processes. Previous BEDASSLE analyses that detected stronger IBE patterns were conducted over much larger geographic scales among much more strongly differentiated populations, such as sky island birds (Manthey and Moyle 2015), lizards occupying a SE Asian archipelago separated by deep ocean trenches (Barley et al. 2015), and a widespread bird occupying much of the Amazon basin (Harvey and Brumfield 2015). These results suggest that the IBE model in BEDASSLE may currently be best suited to populations that are deeply diverged in both time and space.

## Conclusions & Future Research

The results presented here demonstrate for the first time that urbanization is associated with a pervasive reduction in genome-wide variation among animal populations. *Peromyscus* spp. Are increasingly important models for investigating natural variation (Bedford and Hoekstra 2015). Given unchecked urbanization, particularly in the eastern United States, it is likely that many white-footed mouse populations in metropolitan areas have experienced similar declines in standing genetic variation. This study examined only a single urbanization gradient. Replicate analyses on white-footed mice in other metropolitan areas, and studies on additional taxa that may be isolated in urban fragments, are necessary to robustly establish a pattern of declining genome-wide variation in urbanizing landscapes.

Preliminary evidence indicates that some loci have experienced selective sweeps in urban white-footed mice (Harris et al. 2013), but the relative impacts of urbanization on historical demography and natural selection have not been fully disentangled for these populations. Using an expanded transcriptome dataset (Harris et al. 2015), we recently identified signatures of selection in NYC populations using an approach that accounts for historical demographic patterns (Harris & Munshi-South, unpublished manuscript). We identified dozens of candidate loci under selection that are associated with metabolic and immune processes. These patterns may reflect changes in diet and biotic pressures (i.e. disease and inflammation) in cities. Thus, the loss of genome-wide diversity documented here does not necessarily preclude local adaptation to highly altered, stressful urban environments.

Evolutionary potential of urban populations could be improved by restoring connectivity between urban forest patches. Enhanced gene flow between urban forests, as well as between urban and suburban areas, would increase overall genome-wide diversity (although may simultaneously break up local adaptation). Habitat networks that promote gene flow in cities can be constructed from even small gardens and green spaces not explicitly dedicated to biodiversity (Goddard et al. 2010; Vergnes et al. 2012). Microevolutionary processes are rarely integrated into urban conservation due to their perceived complex nature and variation between taxa, but could be useful metrics for assessing landscape connectivity (Stockwell et al. 2003; Kinnison et al. 2007). Converting “gray” infrastructure into “green” networks could also simultaneously address conservation goals while delivering co-benefits to humans in the form of biodiversity experiences (Tanner et al. 2014). A rich literature indicates that humans benefit in myriad ways from access to nature (Fuller et al. 2007), but integrative approaches to urban wildlife conservation are a major area for future growth (Shwartz et al. 2014).

### Data Archiving Statement

The DNA sequencing reads will be deposited in NCBI’s Short-read Archive (SRA). Other data files will be deposited on Dryad.

## List of Supplementary Information

**Table S1.** Correlation coefficients calculated between % impervious surface and human population sizes calculated at buffers around study sites of 500 m, 1 km, 1.5 km, and 2 km.

**Table S2.** Summary population genomic statistics calculated for all nucleotide positions (variant and fixed).

**Figure S1.** Cross-validation (i.e. a-score optimization) to identify the optimal number of principal components to retain for DAPC without overfitting4

**Figure S2.** Compoplot / bar plot result from DAPC analysis on all 23 populations.

**Figure S3.** Scatterplot of first two discriminant functions from DAPC for populations on Long Island.

**Figure S4.** Cross-validation of results from ADMIXTURE for K = 1 – 12.

**Table S3.** Matrix of pairwise F_ST_ (above diagonal) and great-circle geographic distance (km; below diagonal) calculated between all pairs of 23 populations.

**Figure S5.** Cumulative current map from isolation by resistance (IBR) modeling in Circuitscape. Lighter areas represent landscape cells with higher cumulative predicted current (i.e. higher movement). All landscape cells with impervious surface % > 70% were assigned resistance = 100, and all cells with impervious surface < %70 were assigned a resistance equal to their percent impervious surface (e.g. 50% impervious surface results in 50X higher resistance than 1% impervious surface).

